# BiaPy: Accessible deep learning on bioimages

**DOI:** 10.1101/2024.02.03.576026

**Authors:** Daniel Franco-Barranco, Jesús A. Andrés-San Román, Ivan Hidalgo-Cenalmor, Lenka Backová, Aitor González-Marfil, Clément Caporal, Anatole Chessel, Pedro Gómez-Gálvez, Luis M. Escudero, Donglai Wei, Arrate Muñoz-Barrutia, Ignacio Arganda-Carreras

## Abstract

BiaPy is an open-source library and application that streamlines the use of common deep learning approaches for bioimage analysis. Designed to simplify technical complexities, it offers an intuitive interface, zero-code notebooks, and Docker integration, catering to both users and developers. While focused on deep learning workflows for 2D and 3D image data, it enhances performance with multi-GPU capabilities, memory optimization, and scalability for large datasets. Although BiaPy does not encompass all aspects of bioimage analysis, such as visualization and manual annotation tools, it empowers researchers by providing a ready-to-use environment with customizable templates that facilitate sophisticated bioimage analysis workflows.

---

Bioimage analysis is a cornerstone of modern life sciences, powering discoveries and insights derived from biological image data. Deep learning has become an invaluable tool for analyzing microscopy datasets, and its application is increasingly widespread in biomedical research [1]. However, the prerequisite for high-level programming skills has often acted as a barrier, limiting accessibility to researchers without a specific computational background [2, 3].

The rapid evolution of deep learning methods, along with the diverse range of bioimage analysis applications, has created a dynamic landscape that requires researchers to adapt continuously. These applications often involve navigating multiple tools and integrating *workflows* that handle tasks such as image segmentation, object detection, tracking, image classification, and reconstruction. To support this diversity, many solutions now either build upon or integrate deep learning approaches in some form [4–15].

For example, deepImageJ [5] integrates pretrained deep learning models into Fiji [4], leveraging resources like the BioImage Model Zoo [16], which facilitates the sharing and reuse of models across a broad spectrum of workflows. Similarly, tools like ZeroCostDL4Mic [6] and ImJoy [7] simplify access to deep learning workflows through Jupyter notebooks and web interfaces. QuPath [14], Ilastik [10], and Icy [17], all Community Partners of the BioImage Model Zoo, offer additional frameworks for incorporating deep learning approaches into bioimage analysis workflows.

Several tools have been developed to support the creation of pipelines for image classification and segmentation [8–11, 15]. In addition, image classification and object detection workflows can be constructed using some of these tools [9, 10, 14, 15].

A great many tools have been developed to support functionalities such as image classification, segmentation, and object detection [4, 7–11, 14–17]. Some of these tools, like Fiji [4] and Icy [17], provide features for building workflows in a pipeline-based paradigm through plugins such as Jipipe and Protocols, respectively. Others, such as Cellpose [8], CellProfiler [9], Ilastik [10], PlantSeg [11], and QuPath [14], are designed with accessibility in mind and offer user-friendly graphical interfaces that lower the barrier for users without a computational background. Tools like QuPath [14] and Cellpose [8] have further enhanced accessibility by integrating deep learning functionalities into their GUIs, enabling seamless workflows for biologists and researchers. However, some tools, such as InstantDL [15], rely primarily on command-line interfaces, which can present challenges for certain user groups. This variety highlights the diverse approaches taken to meet the needs of different audiences in the bioimage analysis community.

Web-based platforms, such as ZeroCostDL4Mic [6] and ImJoy [7], attempt to mitigate the barriers of software installation, though this comes with trade-offs, such as reduced flexibility. For example, ZeroCostDL4Mic relies on Google Colab, which requires ongoing updates to align with changes on the platform. To improve portability and reproducibility, the same authors introduce [18], which uses Docker containers to encapsulate the notebooks along with their dependencies.

In terms of versatility, tools like deepImageJ [5] are primarily focused on making predictions with pretrained models and do not yet support training on custom datasets, limiting their adaptability for researchers with more specific experimental needs. These constraints can be a drawback for users seeking tools that offer flexibility for customized bioimage analysis setups.

To address specific gaps in the current bioimage analysis landscape, we present BiaPy (https://biapyx.github.io), an open-source library designed to streamline a variety of bioimage analysis tasks, particularly for multi-channel, multi-dimensional microscopy data (Fig. 1a). BiaPy simplifies the analytical process by supporting both traditional convolutional neural networks (CNNs) and modern Transformer architectures [19] within a single unified interface. This mix of established and advanced models allows researchers to apply deep learning consistently, accurately, and flexibly, aligning with the rapidly evolving methods of bioimage analysis.

**Fig. 1:**
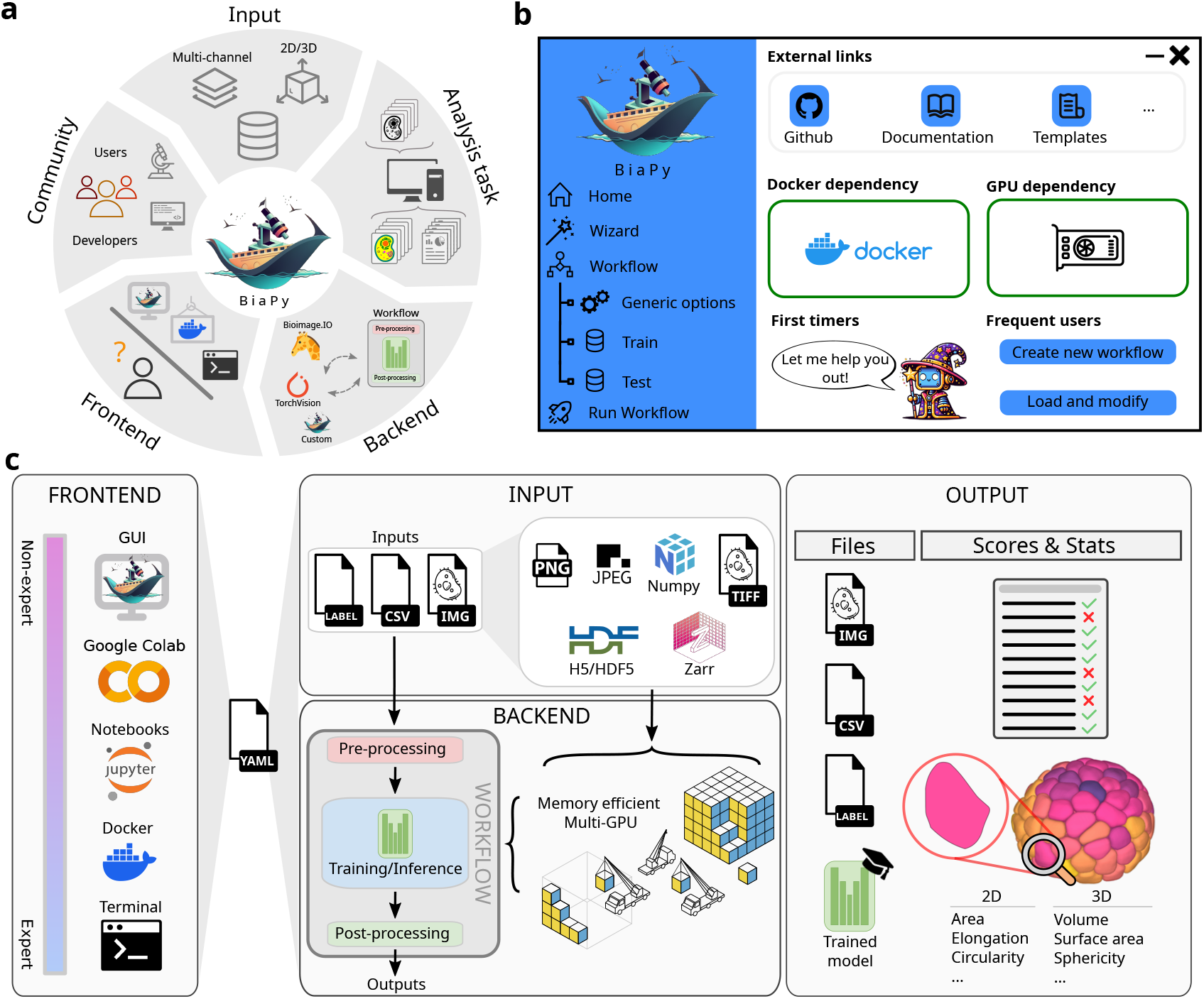
BiaPy environment and scope. **a**. General overview. By addressing their distinct needs, BiaPy accommodates life science users and computer vision developers. Multi-channel, 2D/3D microscopy datasets serve as inputs for BiaPy workflows. Recurrent bioimage analysis tasks are executed by workflows that can be customized, imported/exported from/to the BioImage Model Zoo [16], or leverage pretrained models from TorchVision. The backend allows for customization, while the frontend caters to a broader user base. **b**. Graphical User Interface (GUI): BiaPy’s GUI serves as an intuitive interface for user interaction, featuring a guiding wizard to assist first-time users through the necessary steps and decisions. **c**. Users have the flexibility to install and run BiaPy through different methods tailored to various experience levels. During execution, a YAML configuration file is created, defining input data and providing instructions for the chosen workflow. BiaPy supports classic image formats (TIFF, Numpy) and modern, memory-efficient formats (H5, Zarr). The workflow consists of three key stages: 1) Pre-processing to prepare the input data. 2) Model training or inference, which processes data segment by segment or patch by patch, employing overlapping/padding strategies and supporting multi-GPU configurations. 3) Post-processing, where probability maps are refined to produce the final results. BiaPy outputs are generated in various formats (e.g., images, tables, text files) and dimensions, with evaluation metrics and morphological statistics provided for comparison with ground truth images where available.

BiaPy is designed to serve life scientists needing accessible tools to accelerate their research, as well as advanced users and developers wishing to integrate custom algorithms from the computer vision research community. Unlike several existing tools that heavily rely on command-line interactions, BiaPy offers a user-friendly interface (Fig. 1b) and zero-code online notebooks (Fig. 1c), making it accessible to users with different levels of technical expertise. The platform uses a single Docker container for its installation, in contrast to other tools that require separate containers for each workflow [18]. This design simplifies installation and enables tighter and more efficient integration between workflows. However, this simplicity comes with trade-offs, such as managing a single dependency set for all supported workflows, which may reduce flexibility compared to the one-container-per-workflow approach. While both approaches enhance portability and minimize conflicts with other software, the single-container design prioritizes ease of use and standardization. However, while BiaPy simplifies certain aspects of deep learning workflows compared to traditional methods, considerable effort is still required for tasks such as data preparation, model training, and result interpretation. These steps remain essential to ensure the effectiveness of the tool in bioimage analysis applications. Additionally, BiaPy is intended to integrate with established platforms, such as through a Fiji plugin and its connection with Brainglobe ^1^, offering users increased flexibility to enhance and streamline their existing bioimage analysis workflows.

BiaPy offers advanced capabilities designed for both local computational environments and larger image facility frameworks. It supports multi-GPU setups and handles large file formats like Zarr and H5, offering scalability for processing substantial datasets and performing computationally intensive analyses (Fig. 1c). Furthermore, BiaPy integrates with popular deep learning ecosystems, such as the BioImage Model Zoo [16]. This integration allows users to import, execute, fine-tune, and export pretrained models within BiaPy’s framework, capitalizing on its computational infrastructure (Fig. 1a). Such interoperability enhances BiaPy’s adaptability for a variety of bioimage analysis tasks. The models can be accessed via the command line, zero-code notebooks, or the GUI, which provides guidance for selecting models compatible with the chosen workflow.

BiaPy follows a streamlined workflow structure consisting of three key elements: data preprocessing, model training (which includes microscopy-specific data augmentation methods), or inference, and data post-processing (Fig. 1c). This structure facilitates the prototyping of new workflows by enabling users to easily select pre- and post-processing methods suited to their specific data and tasks. The library also allows users to switch between various state-of-the-art deep learning models available within BiaPy and its compatible projects, such as TorchVision ^2^ and the BioImage Model Zoo. This design simplifies the process for developers, enabling them to focus on individual parts of workflows rather than needing to build entire pipelines from scratch.

BiaPy is built on Python and PyTorch as its backend for deep learning, leveraging an accessible and widely adopted computational environment. It includes memory optimization strategies, such as automatic mixed precision for faster training on GPUs and efficient RAM utilization, making it adaptable to a variety of computational setups and hardware configurations. For more demanding analyses, BiaPy supports multi-GPU configurations, allowing both training and inference tasks to be efficiently distributed across multiple GPUs, reducing computational bottlenecks. Multi-GPU operations are implemented in chunks to accommodate different system capabilities, enhancing flexibility and scalability (see Fig. 1c).

Each BiaPy workflow is encapsulated in a single configuration file (Fig. 1c), which users can modify directly in a standard text editor or generate through the GUI or zero-code notebooks. Sharing workflows is facilitated by sharing this configuration file, along with model weights if needed for inference. BiaPy workflows are designed to process raw images and corresponding labels or annotations, producing outputs such as predicted annotations, analysis reports, or pretrained models, depending on the task at hand. To improve user accessibility, the GUI includes a Wizard feature that guides users through the workflow setup process by asking simple, task-oriented questions, thereby avoiding technical language that may require prior knowledge [20]. The final configuration is determined based on user input, supplemented by default settings to foster optimal performance across a range of scenarios, making it easier for users to generate configuration files without needing advanced technical expertise.

In its current release, BiaPy supports various workflows for both 2D and 3D image data, including instance and semantic segmentation, object detection, image denoising, single-image super-resolution, self-supervised learning, image classification, and image-to-image translation (see Methods). The instance segmentation workflow follows a bottom-up approach (Fig. 2a), where objects of interest in the input image are detected, segmented, and classified. The model learns binary masks, contours, and, optionally, the distance map of the objects. These representations are combined to generate seeds for a marker-controlled watershed transform, which produces individual instances.

**Fig. 2:**
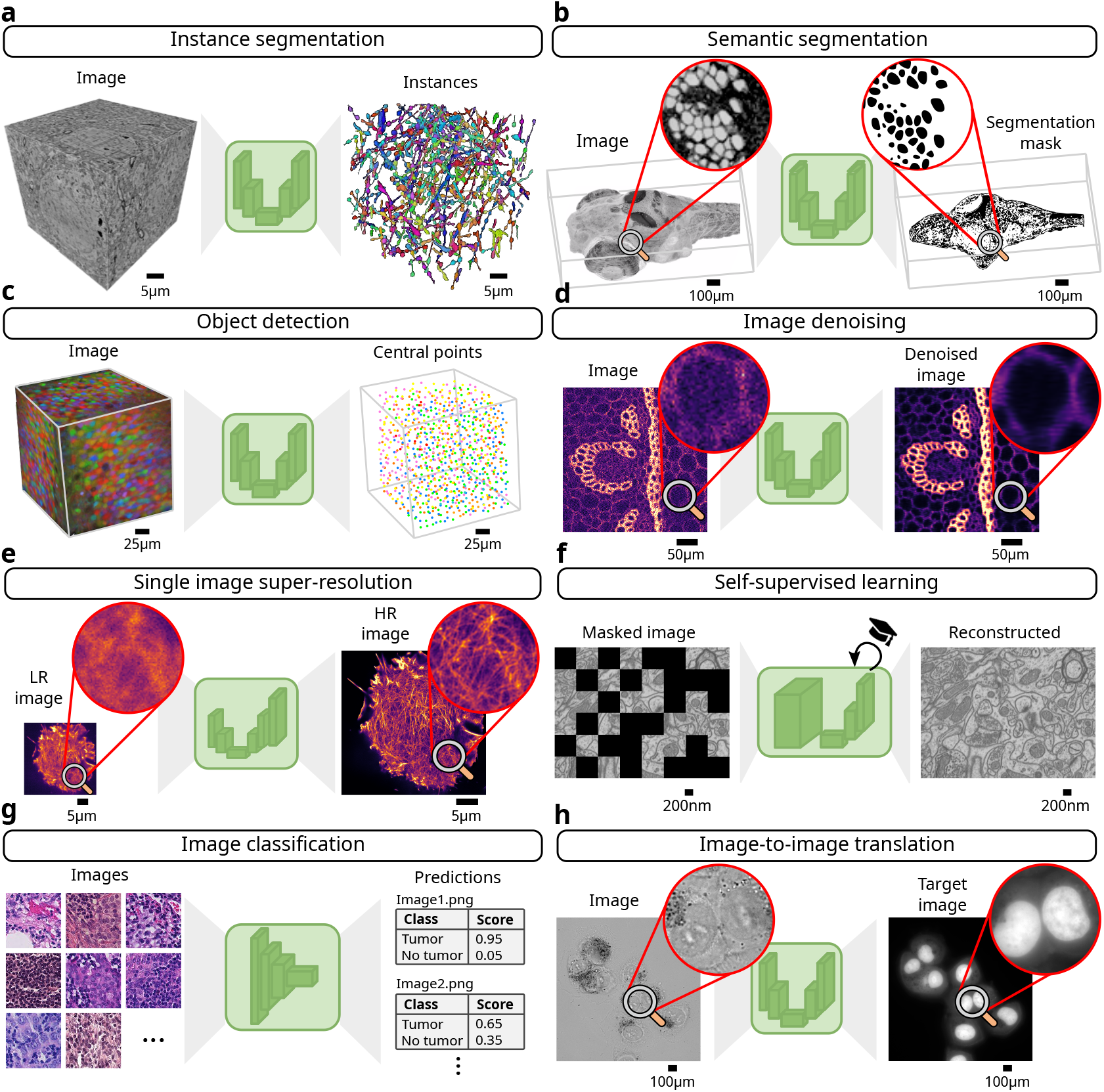
Overview of BiaPy workflows. BiaPy supports the following workflows for 2D/3D grayscale, RGB and multi-channel images: **a**. Instance segmentation involves detecting, segmenting, and classifying individual objects using a bottom-up approach by learning the binary masks, contours, and (optionally) the distance map of the objects of interest. **b**. Semantic segmentation associates a label to every pixel or voxel of the input image. **c**. Object detection localizes objects in the input image, not requiring a pixel-level class, by extracting individual points at their center of mass. **d**. Image denoising removes noise from the input image without needing clean reference data. **e**. Single image super-resolution reconstructs high-resolution (HR) images from low-resolution (LR) ones. **f**. Self-supervised learning pretrains a backbone model on a pretext task without labels, enabling transfer to downstream tasks in labeled datasets. **g**. Image classification labels full input images as belonging to a predefined set of classes. **h**. Image-to-image translation maps input images into target images, e.g., to produce stained images from unstained ones.

In the semantic segmentation workflows (Fig. 2b), each pixel or voxel in the input image is assigned a label that defines its class or category. For object detection (Fig. 2c), BiaPy employs a center-of-mass strategy, commonly used in bioimage analysis, instead of using bounding box annotations.

BiaPy also offers workflows to enhance image quality, such as denoising (Fig. 2d) and super-resolution (Fig. 2e), which produce cleaner or higher-resolution versions of the input images. Additionally, BiaPy includes self-supervised learning workflows (Fig.2f), where models are pretrained on a pretext task without labels. This method allows models to learn a representation that can later be fine-tuned for downstream tasks on smaller, labeled datasets.

Finally, BiaPy provides both image classification workflows (Fig.2g), where models are trained to classify the entire input images into predefined classes, and image-to-image translation workflows (Fig.2h), where input images are mapped to target images, e.g., to convert them to another modality. Further details on these workflows can be found in the Methods section.

In conclusion, BiaPy offers a versatile and portable solution, providing a range of workflows for bioimage analysis that support both 2D and 3D image data. Its combination of an intuitive GUI, zero-code notebooks, and Docker integration makes it accessible to users with different levels of technical expertise. Furthermore, this combination ensures accessibility across a range of hardware configurations, from high-performance machines with GPUs to basic setups without GPU support. BiaPy is designed with a focus on reproducibility and flexibility, making it a valuable tool for both computer vision experts and life scientists. While it does not aim to replace existing platforms, it complements current tools by addressing specific gaps, such as multi-GPU support and adaptability to large datasets. As an open-source initiative, BiaPy encourages collaboration and contributions from the scientific community with the goal of empowering researchers and supporting advancements in bioimage analysis.

Future developments of BiaPy will focus on integrating with established platforms like Fiji, QuPath, napari, CellProfiler, and Icy, enhancing accessibility and usability for biologists. This integration aims to make deep learning workflows more seamless within existing analysis pipelines, addressing the broader challenge of navigating an expanding array of frameworks in bioimage analysis.

## Code availability

The complete source code for the BiaPy platform, encompassing the library’s main code, GUI, and associated documentation, is accessible at https://github.com/BiaPyX. For comprehensive documentation, video tutorials, and use-case examples, please refer to BiaPy’s documentation website (https://biapy.readthedocs.io/en/latest/).

## Acknowledgments

This work is partially supported by grant GIU23/022 (to I.A-C.) funded by the University of the Basque Country (UPV/EHU), grants PID2021-126701OB-I00 (to I.A-C.) and PID2023-152631OB-I00 (to A.M-B.), funded by the Ministerio de Ciencia, Innovación y Universidades, AEI, MCIN/AEI/10.13039/501100011033, and by “ERDF A way of making Europe”. P.G-G. has been funded by Margarita Salas Fellowship – NextGenerationEU. J.A.A-S. work has been funded by the Junta de Andalucía (Consejería de economía, conocimiento, empresas y Universidad) grant PY18-631 co-funded by FEDER funds. L.M.E. also wants to thank the PIE-202120E047-Conexiones-Life network for its networking and input. I.H-C. and A.M-B. received funding from the European Union through the Horizon Europe program (AI4LIFE project with grant agreement 101057970-AI4LIFE). Funded by the European Union. Views and opinions expressed are, however, those of the authors only and do not necessarily reflect those of the European Union. Neither the European Union nor the granting authority can be held responsible for them. I.H-C. also acknowledges the support of the Gulbenkian Foundation (Fundação Calouste Gulbenkian).

## Author contributions

Conceptualization: D.F-B., A.C., L.M.E, D.W., A.M-B. and I.A-C. Software: D.F-B., J.A.A-S., I.H-C., L.B., A.G-M., P.G-G., C.C. and I.A-C. Validation: D.F-B., I.H-C., L.B., A.G-M., P.G-G., C.C. and I.A-C. Writing—initial outline: D.F-B., A.M-B. and I.A-C. Writing—original draft: D.F-B., A.M-B. and I.A-C. Writing—review and editing: all authors. Visualization: D.F-B. and I.A-C. Supervision: L.M.E, D.W., A.M-B. and I.A-C. Funding acquisition: A.C., L.M.E, D.W., A.M-B. and I.A-C.

## Methods

The following sections provide technical descriptions and outline the software requirements for using BiaPy. Given that BiaPy is an actively evolving project, users and developers are encouraged to refer to the official documentation on the BiaPy website (https://biapyx.github.io/) for the most up-to-date information.

### Design goals of BiaPy

The design philosophy of BiaPy addresses the multifaceted challenges inherent in the field of bioimage analysis while democratizing access to cutting-edge computational techniques. At its core, BiaPy strives to fulfill several key objectives:

1. **Unified framework:** BiaPy is crafted as a unified platform, integrating both backend and frontend functionalities. This design ensures accessibility and benefits for both novice and expert users.
2. **Versatility in handling image modalities:** BiaPy is engineered to support a wide range of image modalities, including multi-channel, frequently anisotropic, 2D, and 3D images, addressing the diversity of bioimaging data.
3. **Customizable for various output targets/tasks:** BiaPy’s architecture is designed to be adaptable to different output targets and tasks, offering customizable solutions for tasks such as semantic segmentation, instance segmentation, object detection, denoising, and super-resolution.
4. **Scalability and optimization:** BiaPy focuses on scalability and optimization, incorporating multi-GPU capabilities and memory optimization strategies to handle computationally demanding bioimage analyses with large datasets.
5. **Community-driven and collaborative approach:** BiaPy is envisioned as a collaborative tool, encouraging contributions and feedback from the research community. This open-source nature fosters continuous improvement and innovation in bioimage analysis.

Through these design goals, BiaPy aims to provide an accessible, portable, and flexible tool for the bioimage analysis community, enabling researchers to harness the power of deep learning and other advanced computational methods in their work. Table 1 provides a detailed comparison of BiaPy’s features with those of similar tools currently available in the bioimage analysis ecosystem.

**Table 1:**
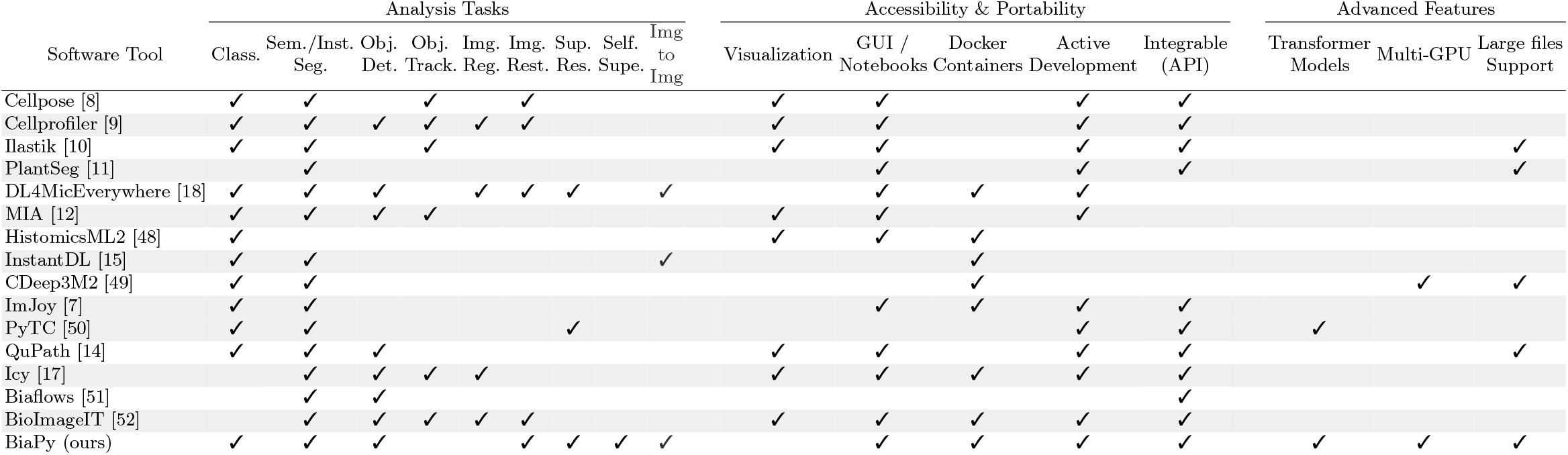
Comparison of features available in BiaPy and similar state-of-the-art software tools. Inclusion in this list is based on (1) target bioimage analysis tasks, (2) accessibility and portability features, and (3) advanced technological features. The terms (‘Class’, ‘Sem./Inst. Seg.’, ‘Obj. Dect.’, ‘Obj. Track.’, ‘Img. Reg.’, ‘Img. Rest.’, ‘Sup. Res.’, ‘Self. Supe.’, ‘Img to Img’) represent Classification, Semantic/Instance segmentation, Object detection, Object tracking, Registration, Image restoration, Super-resolution, Self-supervised learning, and Image-to-Image translation, respectively. Notice the image restoration category includes denoising and deconvolution tasks. The comparison is primarily focused on BiaPy GUI as a non-command-line alternative for operating the software rather than a comprehensive visualization tool. We consider projects as actively developed when their repositories contain 12 or more commits in the last year.

### Installation and usage: from novice to expert

For a comprehensive guide to installation, refer to https://biapy.readthedocs.io/en/latest/get_started/installation.html. BiaPy offers multiple avenues for installation and usage to accommodate users with varying technical proficiencies:

– **Graphical user interface (GUI)**: BiaPy includes a GUI for users seeking a user-friendly interface. The GUI performs checks for Docker image installation and GPU availability, enabling easy workflow execution. Currently, the GUI is designed to operate with a single GPU (no multi-GPU).
– **Google Colab integration**: BiaPy provides code-free notebooks in Google Colab, lowering entry barriers and enabling researchers to use BiaPy without local installations.
– **Jupyter notebooks**: Users can access templates and executable Jupyter notebooks locally for each workflow, accompanied by example datasets for better understanding.
– **Docker**: BiaPy is available as a Docker image, ensuring a consistent environment across systems. Two containers support different PyTorch/CUDA versions for broader accessibility.
– **Command line**: For users familiar with conventional workflows, obtaining BiaPy is as straightforward as executing a repository clone command or using standard package installation tools (e.g., *pip*). Subsequent installation of dependencies grants users direct access to BiaPy’s functionalities.

### Software dependencies and hardware requirements

BiaPy has undergone successful testing on Linux, MacOS, and Windows operating systems. Given the deep-learning core of BiaPy, a machine equipped with a GPU is recommended for optimal training and execution speed. Users accessing BiaPy through its GUI need only Docker, as the GUI operates BiaPy indirectly using Docker. The GUI is provided as a binary file, available for download from BiaPy’s documentation (https://biapy.readthedocs.io/en/latest/get_started/installation.html) or GitHub (https://github.com/BiaPyX/BiaPy-GUI) pages. After downloading, it can be launched with a simple double-click. For other cases, such as Google Colab, BiaPy’s dependencies are installed using *pip*.

To accommodate a broader user base, BiaPy is distributed through two separate Docker containers: one based on PyTorch 2.1 and another on PyTorch version 1.2.1, corresponding to CUDA versions 11.8 and 10.2, respectively. This approach ensures compatibility with older GPUs that may have outdated drivers while retaining full functionality in BiaPy.

### Multi-GPU setting

BiaPy offers training on a multi-GPU setting using PyTorch’s distributed data-parallel (DDP) training, a widely adopted paradigm for single-program multiple-data training. Moreover, BiaPy introduces a novel strategy for multi-GPU inference (depicted in Fig. 1c). Unlike the conventional method of distributing all test images across available GPUs for accelerated processing, BiaPy’s approach is built to accommodate biological microscopy image data at scale, addressing challenges posed by very large images. More specifically, our method addresses the constraints related to memory and disk space. BiaPy enables multi-GPU processing per image by chunking large images into patches with overlap and padding to mitigate artifacts at the edges. Each GPU processes a chunk of the large image, storing the patch in its designated location within an output file, typically in Zarr or H5 format. These file formats facilitate reading and storing data chunks without requiring the entire file to be loaded into memory. Consequently, our approach allows the generation of predictions for large images, overcoming potential memory bottlenecks.

### BiaPy workflows

BiaPy is designed to process multi-channel, frequently anisotropic, 2D, and 3D images by means of a variety of workflows. These workflows are enhanced by the integration of models from the pretrained BioImage Model Zoo [16] and TorchVision models ^3^, accessible via their official repository, when applicable to the specific task. Additionally, BiaPy supports our own custom deep learning models, details of which are provided for each specific task. The following section outlines the technical specifics and the diverse model support of the existing BiaPy workflows:

#### Semantic segmentation

Images are processed to assign a class to each pixel or voxel of the input image. For this task, the deep learning models currently available in BiaPy are custom U-Net [21, 22], Residual U-Net [22, 23], ResUNet++ [24], Attention U-Net [22, 25], MultiResUnet [26], Squeeze-and-Excitation (SE) block U-Net [22] and UNETR [27]. Training image patches can be selected based on a probability map generated from the ground truth masks. This approach ensures a higher frequency of patches containing a foreground class is input into the deep learning model. Apart from the full 3D workflow implemented, at test time, a workflow segmenting 2D images can also produce a 3D output if test images constitute a 3D stack. To address potential 3D inconsistencies and enhance the smoothness of the predictions, BiaPy offers two post-processing methods. These include applying median filtering either along the z-axis or concurrently on the x- and y-axes. The semantic segmentation results are evaluated using the intersection over union (IoU) score.

#### Instance segmentation

The goal of this workflow is to assign a unique identifier, i.e., integer, to each object of interest in the input image. For this task, BiaPy currently supports the following models: U-Net [21, 22], Residual U-Net [22, 23], ResUNet++ [24], Attention U-Net [22, 25], MultiResUnet [26], SE U-Net [22] and UNETR [27]. BiaPy uses a bottom-up approach for instance segmentation, where models are trained to predict intermediate representations of the object masks, such as binary masks, boundaries, central points, and/or distance maps [28, 29]. Then, marker-controlled watershed [30] is used to convert these representations into object instances. In addition to the post-processing methods described for semantic segmentation, predicted instances can be refined using morphological operators or based on a Voronoi tessellation [31]. The quality of the output instances can be measured by a large variety of metrics, including average precision, association, and matching metrics [29].

#### Object detection

This workflow aims to localize objects within the input image without requiring pixel-level classification. Common strategies include producing bounding boxes around objects or pinpointing their center of mass [32], the latter being the approach employed by our tool. Multi-class central points are also supported in this workflow. BiaPy supports several models for this task, such as U-Net [21, 22], Residual U-Net [22, 23], ResUNet++ [24], Attention U-Net [22, 25] and SE U-Net [22]. For training, BiaPy uses additional input CSV files that list the coordinates for the center of mass of each object within each class. Then, those coordinates are used to create a target image with point masks. During inference, the model computes probabilities for each center of mass, which are subsequently processed to identify final detections. BiaPy includes several post-processing methods, such as non-maximum suppression, to remove close points based on a predefined radius. Complete instances can be reconstructed from the predicted centers using marker-controlled watershed [30], offering adaptability to various object shapes. The same filtering techniques applied in the instance segmentation workflow are available here, tailored to the instances’ shapes. For evaluation, BiaPy computes precision, recall, and F1 metrics for the final predicted points based on a predetermined tolerance to assess the model’s accuracy.

#### Image denoising

This workflow aims to eliminate noise from images. Our library incorporates the Noise2Void framework [33], an unsupervised method that only needs as input the noisy images and tries to reduce the probabilistic noise. Available models for this task include U-Net [21, 22], Residual U-Net [22, 23], ResUNet++ [24], Attention U-Net [22, 25] and SE U-Net [22].

#### Single image super-resolution

This workflow focuses on reconstructing high-resolution (HR) images from their low-resolution (LR) counterparts. Available models include the Enhanced Deep Residual Network (EDSR) [34], Deep Residual Channel Attention Networks (RCAN) [35], Deep Fourier Channel Attention Network (DFCAN) [36], Wide Activation for Efficient and Accurate Image Super-Resolution (WDSR) [37], U-Net [21, 22], Residual U-Net [22, 23], ResUNet++ [24], Attention U-Net [22, 25], MultiResUnet[26], SE U-Net [22]. For 3D images, only the custom U-Net-like models are supported. For evaluation, the popular peak signal-to-noise ratio (PSNR) metric is calculated between the reconstructed HR images and the available ground truth data.

#### Self-supervised learning

In this workflow, a backbone model is pretrained by solving a so-called pretext task without the need for labels. This approach allows the model to learn a representation that can later be transferred to solve a downstream task using a labeled, albeit smaller, dataset. In BiaPy, we adopt two pretext tasks: reconstruction and masking. The reconstruction task involves recovering the original image from a degraded version of it [38]. In the masking pretext task, random patches of the input image are masked, and the network is trained to reconstruct the missing pixels or voxels [39]. Available models include the Vision Transformer (ViT) [19], Masked Autoencoder (MAE) [39], EDSR [34], RCAN [35], DFCAN [36], WDSR [37], U-Net [21, 22], Residual U-Net [22, 23], ResUNet++ [24], Attention U-Net [22, 25], MultiResUnet[26], SE U-Net [22] and UNETR [27]. For evaluation purposes, the PSNR metric is calculated, as both pretext tasks aim to recover an image from a distorted input.

#### Image classification

The aim of this workflow is to assign a specific label to the whole input image. The custom models available for this task include a simple convolutional neural network, EfficientNet [40] and ViT [19]. Popular evaluation metrics such as accuracy, precision, recall, and F1 score are calculated to assess the performance of the models.

#### Image-to-image translation

This workflow aims to map input images to corresponding target images, a process commonly referred to as “image-to-image translation”. It can be applied to various tasks, such as image inpainting, colorization, or super-resolution (with a scale factor of 1×). In bioimage analysis, this approach is useful for virtual staining [41, 42], where a model is trained to produce stained images from unstained tissue images or to transfer information between different stains. All models available in BiaPy for single-image super-resolution are compatible with this workflow.

### Microscopy-tailored data augmentation

BiaPy incorporates a range of data augmentation methods commonly used in the classification of natural images but customized for 3D and multi-channel microscopy images of various resolutions. These methods encompass Cutout [43], CutBlur [44], CutMix [45], CutNoise (a variation of CutBlur with additional noise), and GridMask [46], among others. Additionally, BiaPy introduces novel data augmentation techniques specifically designed to mimic typical distortions found in microscope image acquisition, such as misalignment or missing sections (e.g., z-slices). BiaPy also integrates with the imgaug library ^4^, allowing for the implementation of custom augmentation strategies.

### Input and output data management

BiaPy provides two distinct approaches to data management for training, validation, and testing datasets. The first approach involves loading data into memory, which accelerates processing but requires considerable memory allocation. In this approach, the validation dataset can be generated by partitioning the training dataset. The alternative approach is to dynamically load data from the disk as needed, which conserves memory but may result in slower processing times.

For all workflows, except for classification, data loaded into memory is cropped to a specific patch size. This cropping can accommodate predefined overlap and padding. In this context, having images stored with uniform size is not necessary. Nevertheless, when data is dynamically loaded from disk, it should be uniform in size to facilitate consistent patch cropping, which is not always the default condition. To address this, BiaPy includes a random cropping feature, allowing users to ensure consistent patch sizes throughout datasets. In the classification workflows, random cropping is automatically implemented whenever the selected patch size does not correspond with the dimensions of the loaded image.

During the inference phase, each test image can be processed in a full-sized setting or by cropping it to a predetermined patch size. In the latter, this cropping process may also incorporate predetermined overlap and padding, similar to the method applied to training and validation datasets. This technique facilitates the reconstruction of the final image while efficiently minimizing memory usage, thereby ensuring scalability, as detailed in Section 1 of the Online Methods. Furthermore, BiaPy employs test-time augmentation by averaging the predictions from multiple orientations of each patch, specifically creating 8 versions in 2D and 16 in 3D, achieved through multiple 90-degree rotations and mirrored versions of the input images.

### Use case examples

BiaPy has been employed in diverse projects encompassing various image modalities, including electron microscopy (EM), confocal microscopy, microcomputed tomography (uCT), and fluorescence microscopy. The range of object shapes, image resolution, and image contrast in those examples illustrate BiaPy’s adaptability to varied scenarios. The following section briefly describes some representative research projects already published using BiaPy:

#### Mitochondria instance segmentation in large EM volumes ([28, 29])

In a previous work [28], we introduced the MitoEM dataset, a large 3D EM volume including mitochondria of very complex morphology and varying size. This dataset challenged existing instance segmentation methods and motivated the creation of the MitoEM challenge^5^ on 3D instance segmentation of mitochondria in EM images. Together with our report of the findings of the challenge [29], we published our own baseline method (U2D-BC) as a BiaPy workflow, which is fully reproducible using the detailed instructions contained in the tutorials section of BiaPy’s documentation site.

#### Modeling of wound healing in *Drosophila* embryos ([47])

This study compiled a dataset of time-lapse sequences showing *Drosophila* embryos as they recover from a laser-made incision. We approached the modeling of the wound-healing process as a video prediction task, employing a two-stage strategy that combines a vector quantized variational autoencoder with an autoregressive transformer. Our trained model successfully generates realistic videos based on the initial frames of the healing process. In this work, a BiaPy workflow was used for the semantic segmentation of the wounds on each video frame.

#### CartoCell: large-scale analysis of epithelial cysts ([31])

A significant challenge in creating neural networks for cell segmentation is the requirement for labor-intensive manual curation to develop training datasets. CartoCell overcomes this limitation by creating an automated image-analysis pipeline in BiaPy, which efficiently leverages small datasets to generate accurate cell labels in intricate 3D epithelial settings. This workflow enables fast generation of high-quality epithelial reconstructions and, thus, detailed analysis of morphological features. A detailed description and step-by-step tutorial of this workflow are available in BiaPy’s documentation site under “Tutorials”.

#### Analysis of stable deep learning architectures for mitochondria segmentation ([22])

Recent advancements in deep learning models have demonstrated remarkable results in mitochondria segmentation; however, the absence of code and detailed training information frequently impedes their reproducibility. This study adheres to best practices, thoroughly comparing leading-edge architectures and various U-net model adaptations, all implemented as full workflows in BiaPy.

### Limitations of BiaPy

While BiaPy covers a wide range of bioimage analysis tasks (see BiaPy workflows), it does not include functionalities for other common tasks like image visualization, registration, or manual annotation. These tasks are fundamental to many bioimage analysis pipelines and are often required for preprocessing or downstream analysis. For such purposes, users are advised to use other established software platforms.

For example, visualization is not natively supported in BiaPy. To address this, we have collaborated with the Brainglobe project (https://brainglobe.info/), an open-source initiative focused on computational neuroanatomy. In this partnership, BiaPy handles largescale image processing, such as brain-wide cell detection, while Brainglobe’s framework is used for visualizing and further analyzing the results. To aid users in implementing this integration, a detailed tutorial is available, which guides users through the setup and execution of workflows combining BiaPy and Brainglobe: https://biapy.readthedocs.io/en/latest/tutorials/detection/brain_cell_detection.html.

## Footnotes

1 https://brainglobe.info/

2 https://pypi.org/project/torchvision/0.1.9/

3 https://pytorch.org/vision/stable/models.html

4 https://github.com/aleju/imgaug

5 https://mitoem.grand-challenge.org/

